# Transcriptome analysis of growth variation in early juvenile stage sandfish *Holothuria scabra*

**DOI:** 10.1101/2020.09.01.273102

**Authors:** June Feliciano F. Ordoñez, Gihanna Gaye ST. Galindez, Rachel Ravago-Gotanco

**Author notes:** Corresponding author at: *The Marine Science Institute, University of the Philippines Diliman, Velasquez St., Diliman, Quezon City, Philippines 1100*, E-mail address (JFF Ordoñez).

## Abstract

The sandfish *Holothuria scabra* is a high-value tropical sea cucumber species representing a major mariculture prospect across the Indo-Pacific. Advancements in culture technology, rearing, and processing present options for augmenting capture production, stock restoration, and sustainable livelihood activities from hatchery-produced sandfish. Further improvements in mariculture production may be gained from the application of genomic technologies to improve performance traits such as growth. In this study, we performed *de novo* transcriptome assembly and characterization of fast- and slow-growing juvenile *H. scabra* from three Philippine populations. Analyses revealed 66 unigenes that were consistently differentially regulated in fast-growing sandfish and found to be associated with immune response and metabolism. Further, we identified microsatellite and single nucleotide polymorphism markers potentially associated with fast growth. These findings provide insight on potential genomic determinants underlying growth regulation in early juvenile sandfish which will be useful for further functional studies.

**Highlights:**

1. The study explores the genomic basis of growth variation in juvenile sandfish by examining gene expression profiles of fast- and slow-growing early juvenile stages from three hatchery populations using RNA-seq.
2. Sixty-six differentially regulated unigenes potentially related to growth variation are associated with several biological and molecular processes, including carbohydrate binding, extracellular matrix organization, fatty-acid metabolism, and metabolite and solute transport.
3. A large number of potential microsatellite and growth category-associated SNP markers have been identified.

## 1. Introduction

The sandfish *Holothuria scabra* is the highest-valued tropical sea cucumber species. Processed into dried form (*bêche-de-mer* or *trepang*), it is regarded as a luxury food item in Asian markets ^1^. However, the increasing global demand for sea cucumbers has led to unregulated harvesting, intensive commercial extraction, and overall decline of wild stocks and production over the past decade across many fishery areas ^2^, the Philippines included ^3–5^. Advancements in hatchery technology ^6,7^ and rearing in mariculture systems ^8^ represent options for stock restoration and sustainable livelihood activities based on hatchery-produced *H. scabra* ^9,10^.

Sandfish culture practice in the Philippines involves spawning induction and production of larvae in land-based hatcheries, relocation of post-metamorphic juveniles to ocean nursery systems, followed by rearing to marketable size in pond-based or sea-pen grow-out setups ^10,11^. Sandfish are transferred to nursery and grow-out systems upon reaching suitable size and weight. Juveniles can be moved to nursery systems upon reaching lengths > 4 mm (on average 35-40 days post-settlement) and can be transferred to grow-out systems upon reaching > 3 g (typically 30-60 days nursery rearing) ^11^. Consequently, faster-growing juveniles reaching minimum size limits can be can be transferred to ocean nursery and grow-out systems in a shorter period compared to their slower-growing cohorts. Transfer of juveniles to ocean-based nursery systems represents significant reduction in production costs associated with hatchery operations and maintenance and may increase production efficiency with the hatchery potentially accommodating more larval production cycles. Reducing the cost of juvenile production is important for economic viability and to advance sandfish culture to commercial scales.

Growth is a key performance trait of economic importance in aquaculture ^12^. Sea cucumbers exhibit high levels of individual growth variation with coefficient of variation (CV) exceeding 50% ^13,14^. Individual growth variation has been attributed to environmental effects during rearing, with higher stocking densities resulting in increased CVs for two sea cucumber species, *Apostichopus japonicus* ^13,14^ and *H. scabra* ^15^. In *A. japonicus*, while crowding stress has a negative effect on food intake, energy allocation and growth of smaller individuals ^13,14^, genetic factors are still considered to exert significant influence on growth heterogeneity ^13^. Improving culture production systems require a better understanding of the factors affecting the growth of individuals, including genetic variability. ^16,17^. Thus, uncovering genomic determinants for growth performance are of scientific and commercial interest. The advent of next-generation sequencing (NGS) technologies has enabled genome- and transcriptome-wide studies, representing opportunities towards the development of genomic technologies to enhance aquaculture production efficiency and sustainability even for non-model organisms^18^. RNA sequencing technology (RNA-seq) is one of the more powerful high-throughput sequencing approaches to identify and profile candidate genes related to differences in production and performance traits ^19,20^, discover genetic markers for population genetics ^21,22^, and phenotypic variation investigations ^23,24^.

Genetics-based studies on individual growth variation in sea cucumbers are currently limited to *A. japonicus*, based on RNA-seq for comparative analysis of gene expression profiles^25,26^. It remains uncertain, however, whether observations from *A. japonicus* are generally applicable to other sea cucumber species such *H. scabra*. In this study, we performed genome-wide transcriptome analysis of *H. scabra* using RNA-seq to infer genetic mechanisms potentially underlying growth variation in the species. We performed *de novo* assembly and characterized the transcriptome of early juvenile stage *H. scabra* from three different Philippine populations. We also examined differential expression profiles of slow- and fast-growing juveniles and identified potential single nucleotide polymorphism (SNP) markers associated with individual growth variation. The results contribute towards improving our understanding of transcriptome-level regulatory mechanisms underlying individual growth variation in juvenile *H. scabra*. This study contributes genomic resources to enable the development of genome-based technologies for aquaculture and fisheries management through marker-assisted selection, population genetics and adaptation studies in sandfish.

## 2. Materials and methods

### 2.1. Sample Collection

*Holothuria scabra* were sampled at two early life history stages. Juveniles were produced at three hatchery facilities: University of the Philippines - Bolinao Marine Laboratory (BOL), Pangasinan; Palawan Aquaculture Corporation, Coron, Palawan (PAC), and; Alson’s Aquaculture Corporation, Alabel, Saranggani (AAC). The locations of these facilities are shown in Figure 1A. At each hatchery, mass spawning of 40-50 adult sandfish was induced ^27^. Developing larvae were reared in larval tanks for 45 days post-fertilization (Stage 1). Each cohort was then sorted into two growth categories according to body length: (i) fast-growing group (‘shooters’, SHO; with total length (TL) ≥ 3.5 mm) and (ii) slow-growing group (‘stunted’, STU; TL < 2 mm (Figure 1B). All samples from SHO and STU were immediately preserved in RNA*later* (Ambion, Inc., TX, USA) and stored at −20°C until further processing. For Stage 2 juveniles (sand conditioning stage), another cohort was produced and reared for 75 days post-fertilization. The body wall tissues of individuals from SHO (TL ≥30 mm) and STU (TL ≤10 mm) were biopsied, preserved in RNA*later*, and stored at −20 °C until use. Stage 2 samples were only collected for BOL and were only included for the *de novo* assembly.

**Figure 1.**
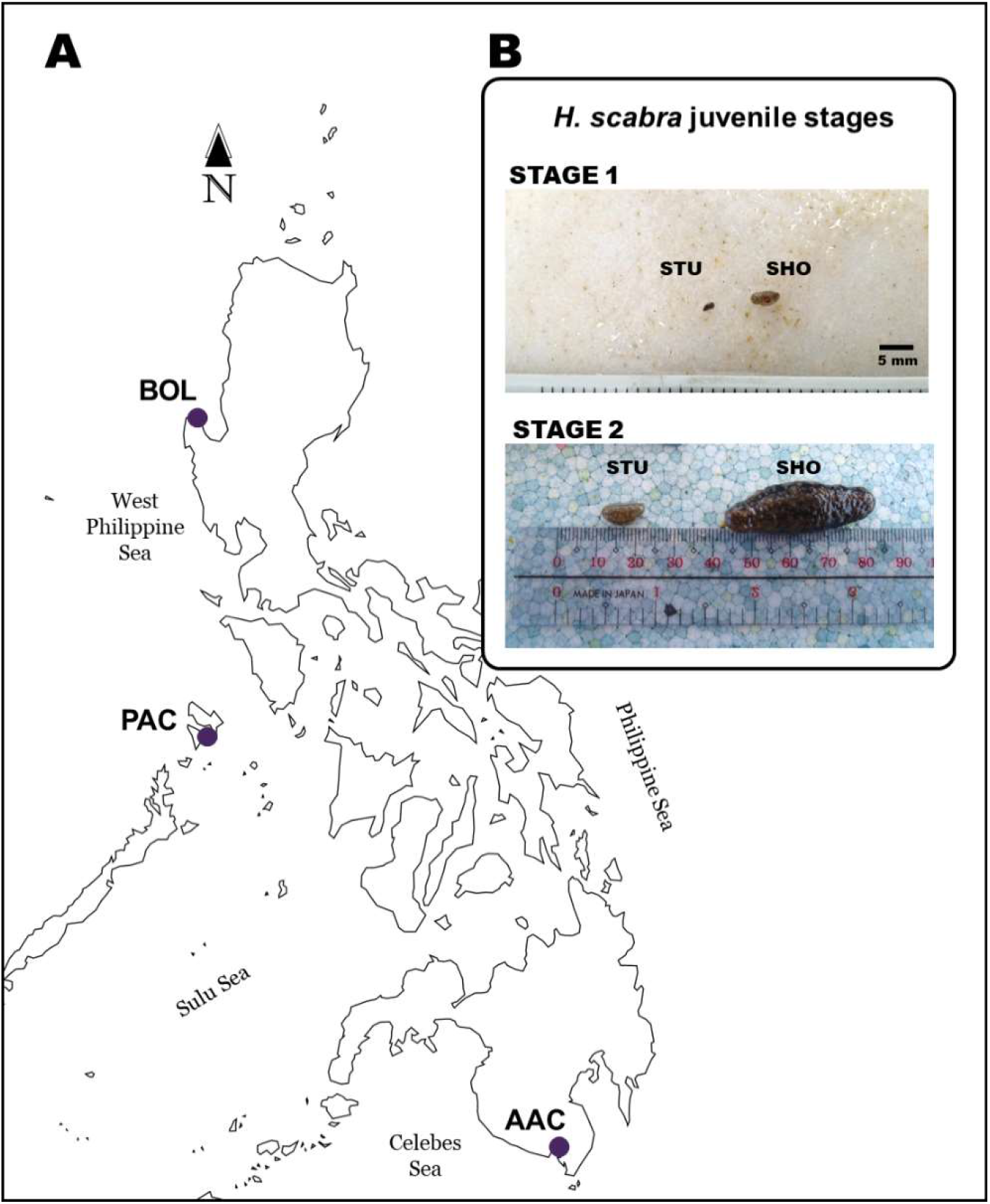
*Holothuria scabra* sample information. (A) A map showing three sandfish hatchery sampling collection sites denoted by purple circles. (B) Sample images of representative individuals from SHO and STU at Stages 1 (45 days post-fertilization) and 2 (75 days post-fertilization).

### 2.2. Total RNA extraction, cDNA library construction, and transcriptome sequencing

Total RNA was extracted from sandfish juveniles using RNeasy Mini Extraction Kit (QIAgen, CA, USA) according to the manufacturer’s instructions. Due to the small size of juveniles, individuals were pooled to ensure recovery of adequate amounts of RNA for sequencing. For Stage 1, each extraction column contained a pool of 7 whole individuals from SHO and 40 whole individuals from STU. For Stage 2, a ratio of 1 SHO: 4 STU was used. RNA quantity and purity were assessed using BioSpec Nano (Shimadzu, Kyoto, Japan) and RNA quality was validated (RNA integrity number > 8) using an Agilent 2100 Bioanalyzer (Agilent Technologies, CA, USA). For Stage 1, biological replicates for each growth category at each of the 3 hatchery populations were prepared for high-throughput sequencing. Stage 2 had no replicates.

cDNA library construction and sequencing were performed by the Beijing Genomics Institute (BGI; Shenzen, China). Library preparation was performed using the Illumina TruSeq™ RNA sample prep kit. A total of sixteen libraries were sequenced on an Illumina HiSeq 2000 (100 bp, paired-end).

### 2.3. Pre-processing, *de novo* assembly, and quality assessment

Initial adapter quality filtering of the raw reads was performed by BGI, which included removal of adapter sequences and reads with ambiguous bases higher than 5%. Further read filtering and trimming was performed using BBDuk from the BBMAP suite v36.11^28^. Reads with overall Q < 20 and with < 70 bp after trimming were further discarded. Error-correction was applied to all reads using Rcorrector v1.0.2^29^. FastQC v0.11.5 ^30^ was used to assess the quality of raw and processed reads.

*De novo* transcriptome assembly was performed using clean reads from all libraries with *in silico* normalization using default Trinity v2.8.4 parameters ^31,32^. To reduce redundancy, contigs from the assembly were clustered using CD-HIT v4.6 with - the following parameters: -s 0.9 -aS 0.9. Transrate v1.0.3 ^33^ was used to filter sequences with low contig scores. Further clustering of potentially related transcripts was carried out using Corset v1.09 ^34^ and salmon v1.1^35^. The longest sequence for each cluster was considered as a “unigene.” Finally, unigenes tagged by Transcriptome Shotgun Assembly (TSA) online submission as contaminants were removed in the final assembly.

Assembly quality and completeness were evaluated using proportion of reads that could be mapped back to transcripts (RMBT), contig ExN50 statistics, Transrate, and BUSCO v3.0.2^36^. RMBT was determined by aligning all clean reads to the final assembly using Bowtie2 v2.2.5 ^37^ and ExN50 was computed using a combination of scripts bundled with Trinity package.

### 2.4. Functional annotation of *H. scabra de novo* transcriptome assembly

Unigenes were queried against various databases and tools capable of predicting potential function of a sequence. Annotation using NCBI non-redundant protein database (nr) was carried out through DIAMOND blast v0.9.29. Unigene annotation was also conducted using Trinotate v3.1.13 (https://trinotate.github.io), which performed sequence homology searching against the SwissProt database ^38^ using blast ^39^, PFAM database ^40^ by HMMER v3.1 ^41^, and association with Gene Ontology (GO) terms ^42^. Trinotate was also used to predict open reading frames (ORFs) by TransDecoder v5.3.0 (http://transdecoder.sourceforge.net), transmembrane region prediction by tmHMM v2 ^43^, signal peptide cleavage site prediction by signal v4 ^44^, respectively. In addition, ORFs were also used to search against the eukaryotic ortholog groups (KOG) using webMGA ^45^ and eggNOG v4.5.1 database using eggNOG-mapper ^46^. Kyoto Encyclopedia of Genes and Genomes (KEGG)^47^ metabolic pathways assignments were performed using the SBH method in the online KEGG Automatic Annotation Server (KAAS)^48^).

### 2.5. Differential expression analysis between SHO and STU

Gene-level differential expression analysis was performed using tximport ^49^ and DESeq2 ^50^. Differential gene expression analysis was only performed on Stage 1 samples due to the lack of replicates for Stage 2. Confounding factors (e.g. batch effects) due to interpopulation variation may not be fully accounted for if DE analysis is performed between SHO and STU across hatchery datasets, which may result in DE inaccuracies. Therefore, DE analysis was performed by comparing SHO against STU for each hatchery dataset. Differential expression of unigenes (DEUs) were considered significant if |log_2_FC| ≥ 2 and an adjusted p-value ^51^ of < 0.01 was observed.

### 2.6. GO and KEGG enrichment analysis of differentially expressed unigenes

GO enrichment analysis of the DEUs was performed using the GOseq ^52^ based on the Wallenius’ noncentral hypergeometric distribution to adjust for gene length bias in the differentially expressed genes. GO terms with corrected p-value < 0.05 were considered significantly enriched. KEGG Pathway enrichment analysis of DEUs was performed using the online tool KOBAS 3.0 ^53^. Reference database for *S. purpuratus* was used as background and hypergeometric test/Fisher’s exact test with FDR-correction ^51^ and a cutoff of < 0.05 was used to test whether identified enriched pathways were significant.

### 2.7. Identification of DNA variants: microsatellites and SNPs

MISA ^54^ was used to identify the potential simple sequence repeats (SSRs) or microsatellite markers in the assembled transcriptome. The parameters were adjusted for identification of at least 10 repeats for perfect mononucleotide motifs, six for dinucleotide, and five for tri-, tetra-, penta-, and hexa-nucleotide motifs.

SNPs discovery was performed using the KisSplice v. 2.4.0-p1 pipeline ^55^. The complete pipeline also allows the evaluation of condition-specificity by testing whether there is a significant association between a SNP and a specific condition (using kissDE v.1.5.0). All programs used in the pipeline were run using default parameters. Only biallelic SNPs were used for downstream analysis.

### 2.8. Hardware and other software used

DE analyses, including DESeq2 and GOseq, were performed using RStudio ^56^, with graphs generated using ggplot2 ^57^, dplyr ^58^, tidyverse ^59^, and pheatmap ^60^. Bioinformatics analyses were performed using either of two local workstations: (i) 6 core Intel^®^ Core(TM) i7-5820K CPU @ 3.30GHz, 4 x 16GB DDR4; and (ii) 2 x 6 core Intel^®^ Xeon^®^ Processor E5-2620 v3 @ 2.4GHz; 8 x16GB DDR4). Both computers run on Ubuntu 16.04.

## 3. Results and Discussion

### 3.1. Sequencing and *de novo* transcriptome assembly for *H. scabra*

To elucidate the genetic basis of growth variation in sea cucumbers, we performed a comparative analysis of gene expression profiles of two growth categories designated as fast-growth (SHO) and slow-growth (STU) in early juvenile stage *H. scabra*. Samples were obtained from three different hatchery populations and sequenced using RNA-Seq.

Over 347 million 100 bp pre-processed reads were obtained from sixteen libraries. Approximately 298 million high-quality paired reads were retained after further trimming, filtering, and error correction (Additional File 1 Table S1) and were used for *de novo* assembly. The initial Trinity assembly generated 369,886 transcripts with a N50 of 1,835 bp, Transrate score of 0.04 (optimal = 0.1), and BUSCO metrics of 94% complete, 5.9% fragmented, and no missing ortholog from the eukaryote database. Reducing the redundancy of the initial assembly resulted in a final assembly consisting of 147,981 unigenes with a N50 of 1,572 bp, average sequence length of 961.1 bp, and a GC content of 38.2 (Table 1).

**Table 1.**
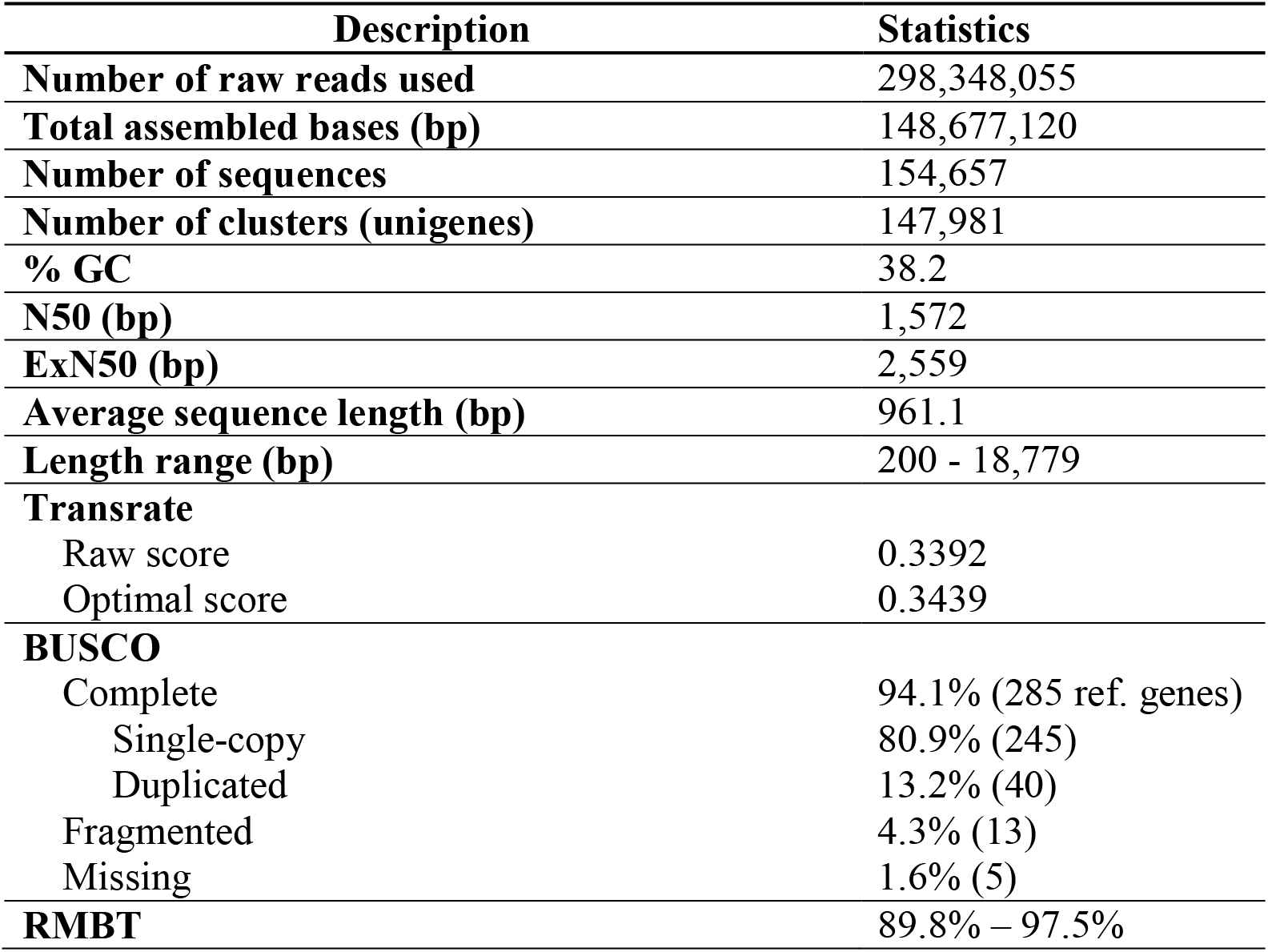
Sequencing and assembly statistics of juvenile *H. scabra* transcriptome.

| Description | Statistics |
| --- | --- |
| **Number of raw reads used** | 298,348,055 |
| **Total assembled bases (bp)** | 148,677,120 |
| **Number of sequences** | 154,657 |
| **Number of clusters (unigenes)** | 147,981 |
| **% GC** | 38.2 |
| **N50 (bp)** | 1,572 |
| **ExN50 (bp)** | 2,559 |
| **Average sequence length (bp)** | 961.1 |
| **Length range (bp)** | 200 - 18,779 |
| **Transrate** |  |
| Raw score | 0.3392 |
| Optimal score | 0.3439 |
| **BUSCO** |  |
| Complete | 94.1% (285 ref. genes) |
| Single-copy | 80.9% (245) |
| Duplicated | 13.2% (40) |
| Fragmented | 4.3% (13) |
| Missing | 1.6% (5) |
| **RMBT** | 89.8% – 97.5% |

Assembly quality was further evaluated using different approaches. Transrate, which estimates the overall quality of the assembly based on the original reads, revealed an assembly score of 0.339 for the *H. scabra* transcriptome, a score higher than the generally acceptable score of 0.22 ^33^. Transcriptome completeness scores using BUSCO showed that the final assembly was 94.1% complete and 4.3% fragmented. Our assembly exhibited low levels of missing single-copy orthologs (1.6% missing), indicating good coverage and quality of the assembly.

To further evaluate the quality of the *de novo* assembly, RMBT and Nx metrics were also calculated. The juvenile sandfish assembly showed a RMBT range of 89.8% - 97.5% and a contig N50 of 1,572 bp. Additionally, ExN50 was calculated as it has been suggested to be more informative than the contig N50, and therefore a more reliable measure of transcriptome assembly quality ^61^. Our assembly showed peak saturation point at 78% of the normalized expression data (E78N50), corresponding to a contig length of 2,559 bp (Additional File 2 Figure S1). Higher quality transcriptome assemblies, however, are expected to produce N50 peak of ~90% (E90N50) of the total expression data ^61^. Lower than E90N50 may indicate that more reads are needed for the assembly. Nonetheless, considering the other quality evaluation metrics used (Transrate, BUSCO, and RMBT), we still assessed the reference assembly to be of good quality and suitable for transcriptome analyses, including marker discovery and differential gene expression analysis.

### 3.2. *H. scabra* transcriptome assembly annotation

Unigenes were translated into proteins using Transdecoder, which predicted 26,124 sequences potentially containing coding regions of at least 100 amino acids in length. In total, 25,761 unigenes (16.7% of the total sequences) were assigned with significant annotations from at least one of the seven query databases (Table 2). The highest number of unigenes with significant hits was reported from nr (16.2%), followed by SwissProt (11.3%), GO (11.5%), PFAM (9.8%), and eggNOG (9.1%). Focusing on unigenes with predicted coding regions, a total of 81.1% (21,195 unigenes) had a significant annotation in one of the query databases. The species distribution from blasting the assembly against nr is shown in Figure 2A. Among the top 15 most represented species, the majority of hits belonged to another holothuroid (*A. japonicus*, 20,344 unigenes), followed by the purple sea urchin *S. purpuratus* (Class Echinoidea; 2,236), and crown-of-thorns *Acanthaster planci* (Class Asteroidea; 1,930).

**Table 2.** Annotation of the juvenile *H. scabra* transcriptome assembly.

|  | **No. of unigenes** | **Proportion (%)** |
| --- | --- | --- |
| **Unigenes** | 154,274 | 100 |
| *Annotation tool* |  |  |
| **nr** | 25,058 | 16.2 |
| **SwissProt** | 17,476 | 11.3 |
| **KEGG** | 13,173 | 8.5 |
| **KOG** | 5,625 | 3.6 |
| **GO** | 17,764 | 11.5 |
| **eggNOG** | 14,100 | 9.1 |
| **PFAM** | 15,176 | 9.8 |
| **SignalP** | 2,415 | 1.6 |
| **tmHMM** | 5,432 | 3.5 |

**Figure 2.**
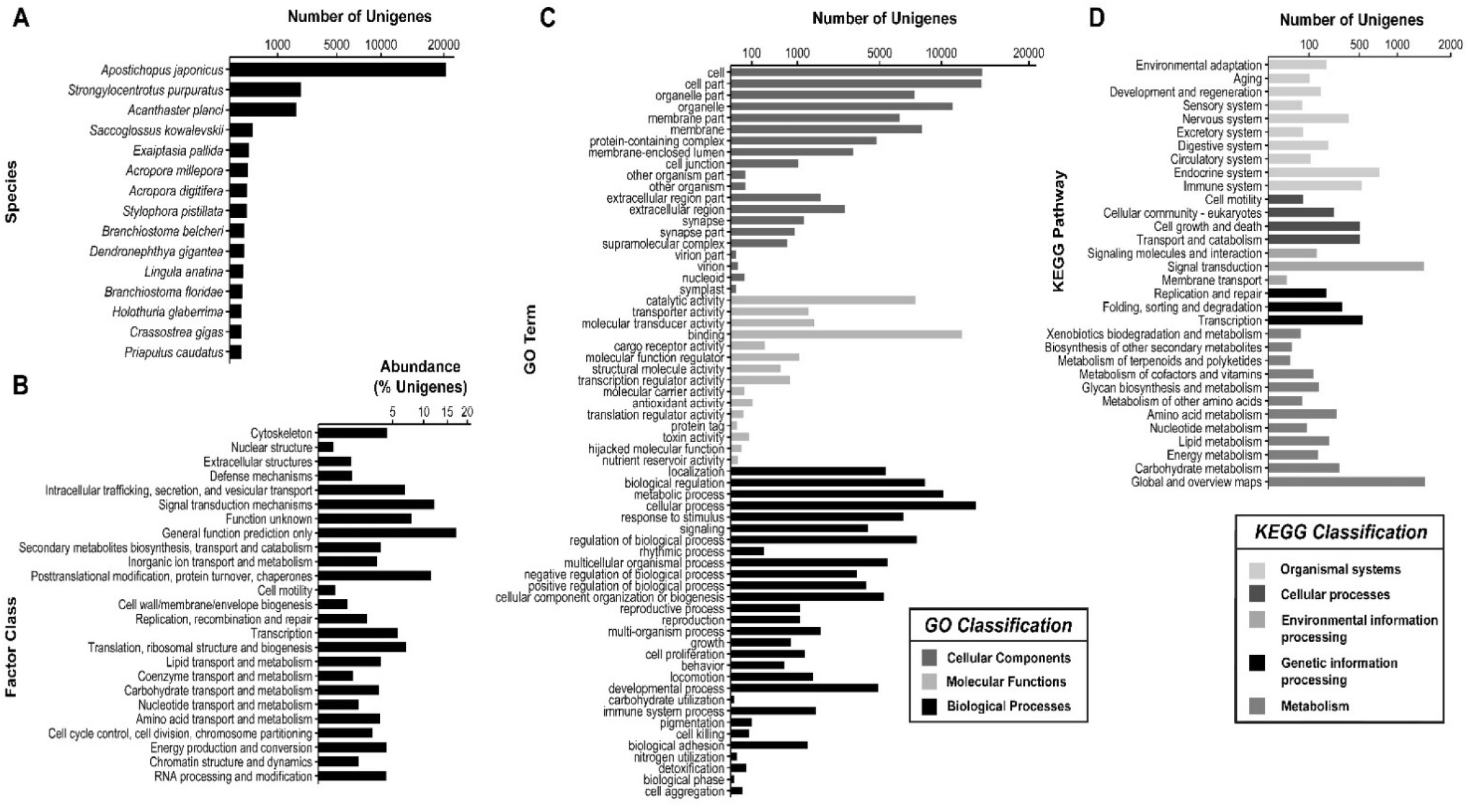
Summary of *H.scabra* transcriptome annotation classified according to top species distribution from nr, eukaryotic <u>ortholog</u> groups (KOG), Gene Ontology (GO), and Kyoto Encyclopedia of Genes and Genomes (KEGG) databases. (A) Top 15 most represented species based on homology search against nr. (B) Frequency distribution of unigenes according to 25 functional categories of KOG. (C) Gene ontology distribution of assembled unigenes for the three general GO classifications. (D) Pathway classification and distribution of unigenes according to five major KEGG categories.

Unannotated unigenes could be attributed to lack of genomic data in public databases for *H. scabra*, misassembled transcripts or chimeras, non-coding (nc) RNAs, and mRNAs that are potentially novel and holothurian-specific^62^. Notably, at least 1,200 sequences in the assembly contain complete protein sequences, ≥ 100 residues in length, and ≥ 10 supporting reads but showed no homology with any genes in the databases used for annotation (data not shown).

To identify and characterize the corresponding functions of the assembled *H. scabra* transcriptome, unigenes were queried against the GO and KOG databases. A total of 5,625 unigenes with predicted ORFs were assigned to one or more KOG annotations (Figure 2B). Among the KOG categories, the “general function prediction only” comprised the largest proportion (17.2% of unigenes with KOG hits), followed by “signal transduction mechanisms” (12.1%). For GO-based annotation (level 2), a total of 17,764 unigenes was mapped to at least one GO term (Figure 2C). Of these, 15,132 unigenes were assigned to Biological Processes (BP), 16,094 to Cellular Components (CC) and 15,538 to Molecular Function (MF). Within the BP category, “cellular process” (13,550 unigenes) and “metabolic process” (10,207) sub-categories were the most represented, while “cell” (14,221) and “cell part” (14,198) were the predominant sub-categories under CC, and “binding” (12,030) and “catalytic activity” (7,678) for MF. Moreover, genes tagged under the term “regulation of growth” (GO:0040008) were also identified, which included sodium- and chloride-dependent GABA transporter 1, nipped-B-like protein A, and signal transducers and activators of transcription 5B (for the complete list, see Additional File 1 Table S2). For characterization of the active biological pathways, unigenes were also queried against KEGG Orthology database. A total of 13,173 unigenes were annotated to 391 KEGG pathways and were classified into 34 pathway categories (Figure 2D). The highest number of hits was identified under the general term “global and overview maps” with 1,477 unigenes with successful hits, followed by “signal transduction” (1,453) and “endocrine system” (743). Using *S. purpuratus* pathway maps as reference for KEGG analysis, 127 metabolic pathways were recovered (Additional File 1 Table S3). The most represented pathway was “metabolic pathways” (1,827 unigenes), followed by “neuroactive ligand-receptor interaction” (332), and “endocytosis” (209).

### 3.3. Gene expression profile comparison of SHO and STU sandfish juveniles

Our results revealed different DEU profiles for the three datasets representing each of the hatcheries. DESeq2 recovered the greatest number of DEUs in AAC (1,324), followed by BOL (831), and PAC (408) (Figure 3A). Differences in DEU profiles across hatcheries may be due to varying physico-chemical conditions during rearing (e.g. temperature, water quality) in different geographic regions. Inherent genetic variation among samples from different biogeographic regions also likely account for DEU profile differences. A population genetic study on *H. scabra* reports genetic divergence among populations of sandfish representing the major marine biogeographic regions in the Philippines ^63^.

**Figure 3.**
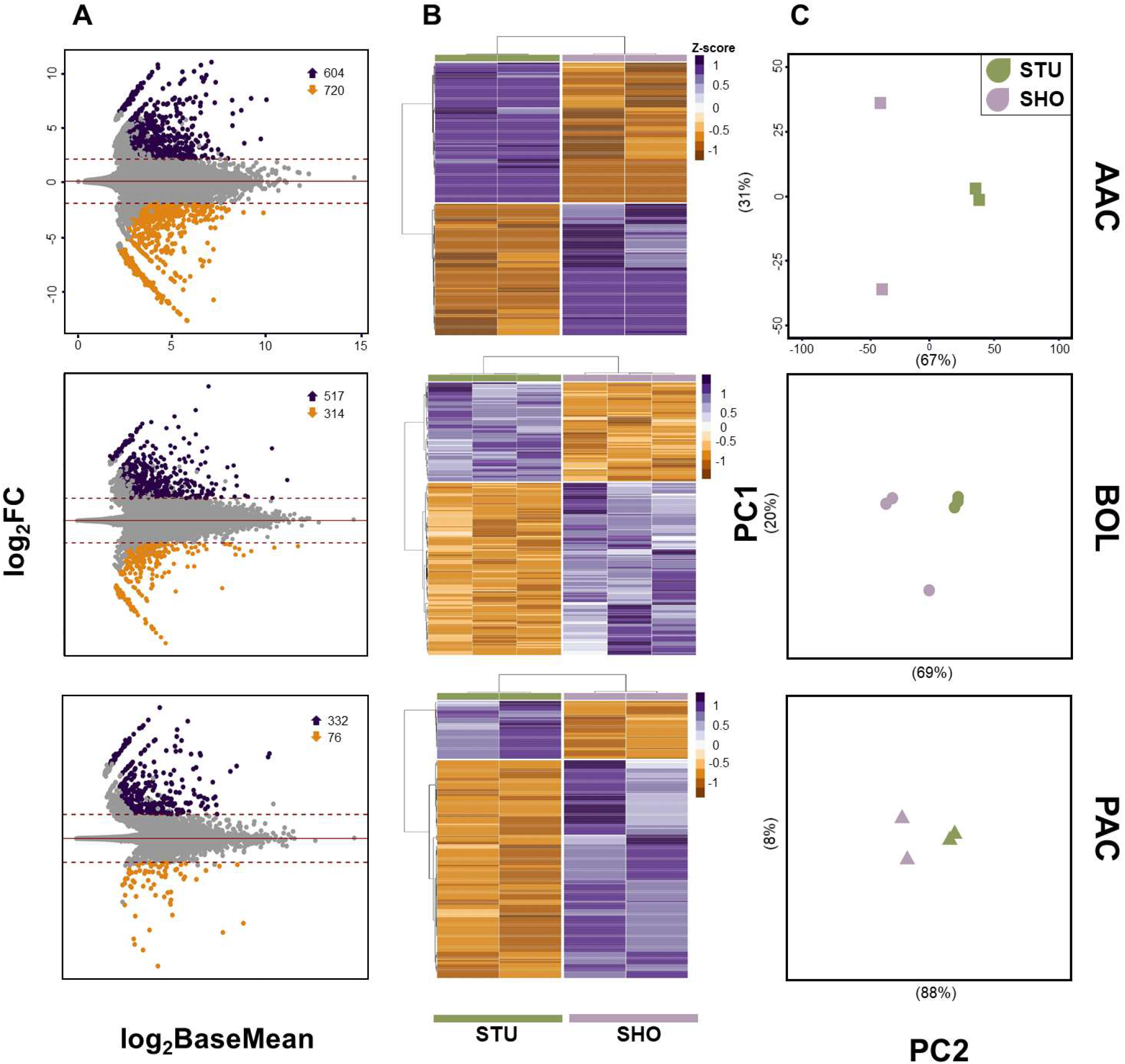
Different representations of gene expression analyses between SHO and STU categories of *H. scabra* for hatchery populations AAC, BOL, and PAC. (A) MA plots highlighting the significant unigenes (FDR p < 0.01) with expression levels of |log2FC| > 2 (denoted by dashed lines). Dots in purple and orange denote upregulated and downregulated unigenes, respectively. Number of upregulated and downregulated unigenes in each hatchery dataset are denoted by up and down arrow, respectively. (B) and (C) are growth-category clustering profiles based on rlog-transformed unigene expression. (B) Heatmaps showing the clustering of SHO and STU samples per dataset. For representation purposes, only the top 200 significant DEUs (log2FC| > 2, FDR < 0.01) were shown. (C) Clustering of the global gene expression in three populations using principal components analysis (PCA).

All three populations shared 66 DEUs that exhibited consistent expression patterns, where 45 and 19 were upregulated and downregulated, respectively (Additional File 2 Figure S2). Of the 66 DEUs, 30 unigenes were assigned with significant (eval: < 1E^−10^) nr annotation (Table 3), while the remaining 36 had no significant hits and potentially encode long non-coding RNA (Additional File 1 Table S4).

**Table 3.**
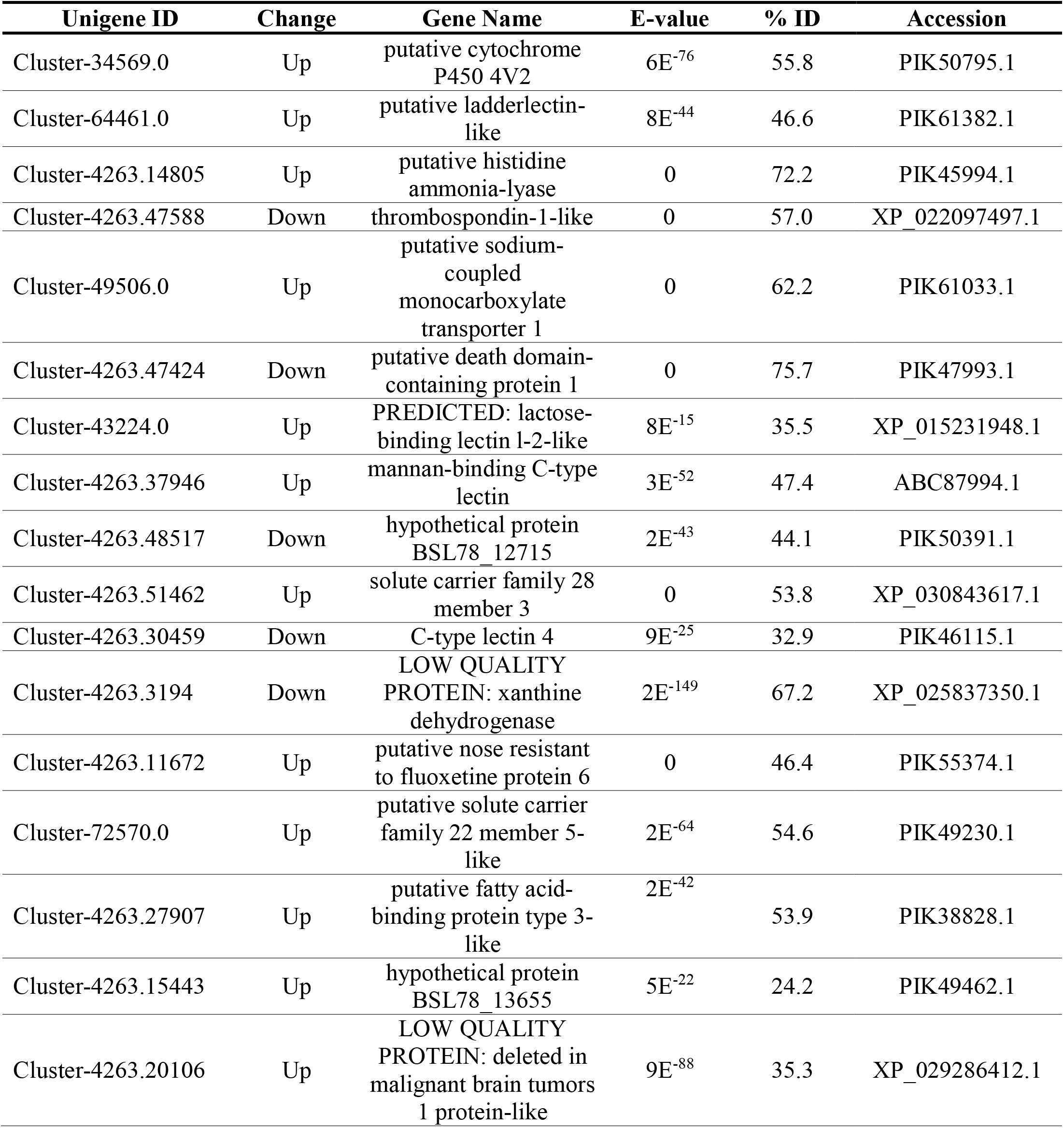
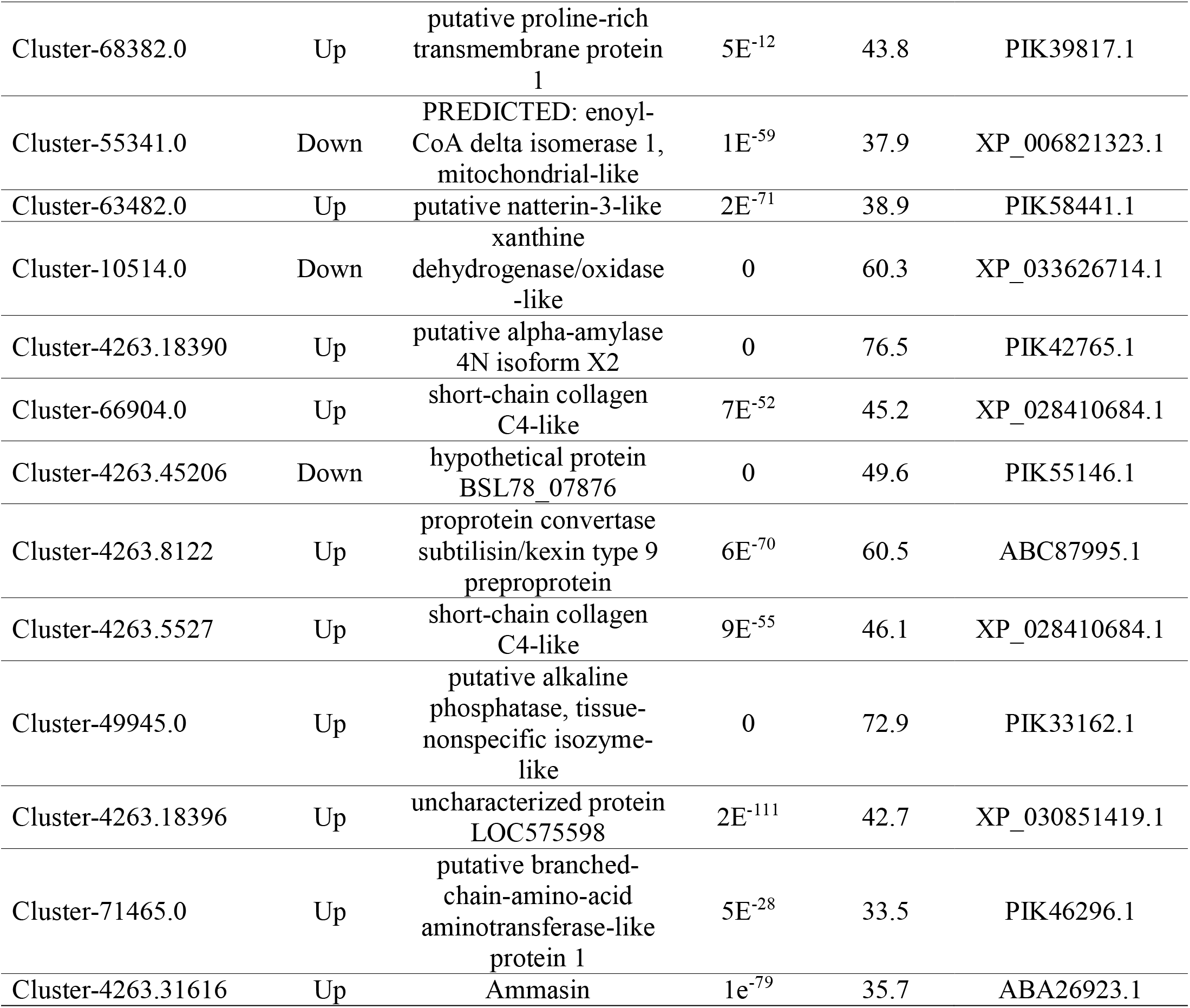
Summary of 30 key differentially expressed unigenes (DEUs) between growth variants (SHO and STU) of juvenile *H. scabra*.

To provide a general overview of the main functions of the identified DEUs, we also performed GO and KEGG analyses on each hatchery dataset. GO terms associated with the DEUs in all datasets were dominated by “cell,” “cell part,” and “membrane,” for CC, and “catalytic activity” and “binding” for MF (Additional File Table S5). GO terms “metabolic process,” “cellular process,” and “biological regulation” were among the most represented GO terms under BP. DEUs in each dataset were observed to be involved in several KEGG pathways but were generally assigned to the following sub-pathways: “global and overview maps,” “lipid metabolism,” “digestive system,” and “transport and catabolism” (Additional File Table S6).

### 3.4. Enrichment analyses of differentially expressed unigenes

Considering that DEU profiles differ considerably among hatchery datasets, we focus on unigenes, enriched GO terms and KEGG pathways that were concordant across hatcheries (AAC, BOL, and PAC); these provide stronger evidence for differential growth and a more robust biological signal of genes and related functions associated with growth variation in sandfish juveniles. Thus we focus our discussion on 30 DEUs that showed consistent expression patterns across all three populations, identified as “key DEUs” (Table 3). We also considered as significant those DEUs that are common between two populations and functionally related to the key DEUs (Additional File 1 Table S7).

#### 3.4.1. GO enrichment analysis

GO enrichment analysis using GOSeq showed highest number of significant (FDR < 0.05) enriched GO terms in AAC (51 terms), followed by PAC (26), and BOL (2) (Additional File 1 Table S8). Enriched GO terms observed in all populations were only related to “carbohydrate binding” (GO:0030246) and “extracellular region” (GO:0005576).

##### 3.4.1.1. DEUs associated with carbohydrate binding

Four key DEUs were enriched in “carbohydrate binding”: lactose-binding lectin l-2-like), C-type lectin 4-like, mannan-binding C-type lectin, and ladderlectin. Notably, these were annotated as genes with C-type lectin-like domains (CTLDs) and, except for the putative C-type lectin 4-like, were all upregulated in SHO. Interestingly, other CTLD-type DEUs were also upregulated in two populations, including putative L-rhamnose binding lectin and ficolin. CTLD proteins are calcium-dependent pattern-recognition receptors (PRRs) that can recognize and bind to carbohydrate moieties (microbe-associated molecular patterns, MAMPs) on microorganisms and activate several immune responses to eliminate pathogens, including the complement pathway, agglutination and immobilization, opsonization, phagocytosis, and lytic cytotoxicity ^64,65^. SHO samples showed upregulation of CTLD genes, which suggests enhanced immune response compared with STU. An intriguing possibility is that immune response to possible pathogen invasion in SHO primarily involves lectin-mediated antimicrobial activities, with CTLD proteins potentially acting as signal receptors, opsonins, agglutinins, or direct antimicrobial effectors. However, not all CTLD genes are deregulated during infection ^65,66^. Therefore, whether differential expression of these immune-related genes is an exclusive consequence of pathogen-dependent immune response remains unclear. Interestingly, we also detected amassin, an upregulated key DEU involved in defense and immunity of echinoderms ^67^, together with several upregulated DEUs common in two populations that are immune-related, including macrophage mannose receptor 1-like, sushi, von Willebrand factor type A, and IgGFc-binding protein. Induced activity of these genes suggests immune response in SHO is highly activated.

##### 3.4.1.2. DEUs associated with extracellular region

The GO term “extracellular region” was also enriched across all populations. Key DEUs identified under this term were ladderlectin, natterin-3, deleted in malignant brain tumors 1 protein (*DMBT1*), short-chain collagen C4 (*CAS4*), proprotein convertase subtilisin/kexin type 9 (*PCSK9*), and thrombospondin-1 (*TSP1*).

The connective tissue of echinoderms comprises extracellular matrix (ECM) proteins, dominated by collagens, proteoglycans, and fibrillin microfibrils ^68^. Proteolytic activities on ECM components are activated to allow ECM transformation and remodeling during pivotal developmental processes, such as morphogenesis, organ development, autotomy, and regeneration ^68–70^. In SHO, we identified upregulated genes involved in ECM modification, which may suggest higher ECM remodeling rate, possibly as a result of faster tissue and organ growth and development. Two key DEUs matched to *CAS4*, which encodes a variant of collagen IV ^71^. Little is known on the role of spongin-related proteins in ECM of echinoderms, but they are assumed to have potentially similar function to collagen IV, including involvement in cell-matrix adhesion, intercellular cohesion, and organismal organization ^72^. In addition, we identified an upregulated DEU homologous with *PCSK9*, an extracellular serine protease that generally performs proteolytic degradation of structural components of ECM (e.g. collagen) to facilitate remodeling of the connective tissue of different organs ^68,73^. Furthermore, DEUs related to ECM-related proteins were identified in two populations, including fibrillin-1, fibropellin-1-like, N-acetylgalactosamine-6-sulfatase, and several serine-type proteases such as *PCSK9* homologs, cuticle degrading serine protease, serine proteinase, chymotrypsinogen-A, and tolloid-like protein. Consequently, these differentially expressed ECM-associated genes potentially play roles in the growth variation in juvenile sandfish by regulating ECM and connective tissue modification.

The key DEU ladderlectin, a gene encoding an extracellular CTLD protein has been suggested to be vital in pathogen clearance because of its ability to opsonize bacteria and viruses ^74,75^. Although the function of ladderlectins in marine invertebrates remains underexplored, it is possible that observed upregulation confers enhanced immunity in SHO, as reported in fish species ^74,76^. A *DMBT1-like* gene was also differentially expressed between STU and SHO. Sandfish *DMBT1* contains the canonical domains CUB, SRCR, and zona pellucida, which have been implicated in the mediation of protein-protein interactions ^77^. *DMBT1* has been suggested to be involved in host disease susceptibility and resistance ^78,79^ and in different developmental processes ^80,81^. A natterin-like gene sharing similar domains (i.e., functionally uncharacterized DUF3421 superfamily and an aerolysin-like pore-forming domain) with naquin (*Thalassophryne nattereri*) natterins was also found to be differentially expressed in all hatchery datasets ^82^. We also identified a DEU upregulated in AAC and PAC that is homologous with natterin-3 of *A. japonicus*. It has been shown that proteins encoded by natterin-like genes can bind and degrade type I and IV collagen and has the ability to destroy pathogens through pore-like complex formation on the target cells, which eventually undergo lysis ^83^. It is plausible that upregulation of these natterin-like genes in SHO influences growth through immune-related mechanisms.

Of the unigenes associated with “extracellular region” that are common across all hatchery datasets, only *TSP1* was upregulated in STU compared to SHO. TSP1 is a trimeric matricellular glycoprotein that has been associated with a wide range of biological functions, including cell adhesion, cell growth, and modulation of cell-to-cell signaling and cell-ECM interactions ^84^. The upregulation of *H. scabra TSP1* in STU is likely to exert an inhibitory effect on growth and development by suppressing the activity of TSP1 targets that regulate growth-related biological activities, such as cellular receptors (e.g. VEGF receptor ^85^) and ECM molecules (e.g. MMPs ^86^).

#### 3.4.2. KEGG enrichment analysis

KEGG enrichment analysis revealed the highest number of significantly enriched pathways (FDR p < 0.05) in AAC with 11 identified KEGG pathways, followed by the BOL and PAC datasets with ten pathways each (Additional File 1 Table S9). KOBAS identified three enriched KEGG pathways common to all populations, namely, “metabolic pathways,” “retinol metabolism,” and “phagosome.”

##### 3.4.2.1. DEUs associated with metabolic pathways

Metabolic pathways (spu01100) comprises several subpathways, including carbohydrate metabolism and energy metabolism. In metabolic pathways, we identified three key DEUs present in all populations, namely, alpha-amylase 2B (*AMY2B*), histidine ammonia-lyase (*HAL*), and alkaline phosphatase (*ALP*).

*AMY2B* encodes for an enzyme that catalyzes the first step in the digestion of dietary starch and glycogen, and thus plays an important role in digestion and energy metabolism. In addition, a DEU similar to sucrase-isomaltase, intestinal-like (*SI*), which is another carbohydrate-degrading enzyme, was identified in the BOL and PAC datasets. Many digestive enzymes, including AMY2B and SI, are endogenous in origin ^87,88^ and their activity can be modulated based on the substrate availability ^89,90^. Consequently, upregulation of *AMY2B* and *SI* in SHO could be a result of increased dietary carbohydrate intake, probably to support the energetically costly metabolic processes concomitant with growth. Growth rate, food intake, and food conversion efficiency has been shown to be generally higher in larger *A. japonicus* individuals compared to their smaller cohorts ^91^.

*HAL* is a gene encoding for an enzyme that catalyzes the first reaction in histidine degradation to urocanic acid and ammonia ^92^. In murine models, high-protein diet has been shown to increase *HAL* expression and concomitantly lower the histidine serum concentrations while undernutrition has been shown to reduce *HAL* activity and decreased overall growth as a consequence of preventing degradation of amino acids, such as histidine, under a condition of dietary protein limitation ^93,94^. Thus, we speculate that lower *HAL* expression in STU compared with SHO may be a consequence lower feeding rate in slow-growing individuals. We also identified a DEU (Cluster-4263.3269) homologous with histamine N-methyltransferase-like (*HNMT*), which exhibited lower expression in SHO compared with STU in AAC and PAC. *HNMT* encodes for an enzyme that catabolizes histamine to 1-methylhistamine ^95^. While speculative, it is possible that higher availability of the histamine-precursor histidine, due to lower *HAL* expression in STU, allows elevated histamine concentration, subsequently causing the upregulation of histaminase *HNMT*. Histamine suppresses feeding in rats in high levels ^96^ and has also been suggested to play a role in feeding behavior of sea cucumber *Leptosynapta clarki*^97^. Nonetheless, the potential associations between *HAL*, *HNMT*, histamine activity, and feeding and growth variation of juvenile sea cucumbers should be experimentally validated in the future.

The final key DEU in metabolic pathways is *ALP*, which encodes the enzyme tissue nonspecific alkaline phosphatase. ALP hydrolyzes a broad class of phosphate monoesters and functions as transphosphorylase in an alkaline environment ^98^. ALP in echinoderms is suggested to play pivotal roles in multiple biological processes, including cell division and differentiation associated with wound healing, mineralization, initiation of regeneration processes, and immune response ^99,100^. Thus, ALP in sandfish may influence growth through its involvement in immunity and morphological development. Interestingly, starvation in *A. japonicus* during periods of inactivation elicits a decrease in ALP levels in the body wall and coelomic fluid of the sea cucumber ^101^, suggesting that ALP activity is also influenced by diet.

##### 3.4.2.3. DEUs associated with Retinol metabolism and Phagosome

Retinol metabolism (spu00830) is another KEGG pathway enriched in all hatchery datasets, which is only represented by putative dehydrogenase/reductase SDR family member 4 (*DHRS4*). DHRS4 is a carbonyl reducing enzyme that participates in the metabolism of endogenous signal molecules, such as retinoic acid ^102^, and in the defense against oxidative stress through detoxification of endogenous lipid-derived aldehydes ^103^. *TSP1* and Actin were enriched in the pathway “Phagosome” (spu04145). *TSP1* exhibited lower expression in SHO compared to STU, which suggests minimal activation of TSP1-mediated pathways, including phagocytosis, in faster-growing sandfish. Actin was upregulated in SHO group of AAC and PAC and may play a role in the regulation of processes that affect growth, including cytokinesis, cell migration, and cell growth ^104,105^.

### 3.5. Other genes potentially associated with growth variation

There were other key DEUs that were not associated with any of the significantly enriched GO and KEGG pathways but may play a role in growth variation in juvenile sandfish.

#### 3.5.1. DEUs associated with purine metabolism

Genes involved in purine metabolism were differentially expressed. Xanthine dehydrogenase/oxidase (*XDH/XOD*) was represented by two different but highly related (aa similarity: 59.5%) key DEUs. *XDH/XOD* catalyzes the terminal step of purine metabolism, converting purine metabolite hypoxanthine to xanthine and subsequently to uric acid ^106^. In addition, we found a DEU in AAC and PAC that is homologous with 5’-nucleotidase (*5NTD*), an enzyme catalyzing the initial step of purine nucleotide degradation (hydrolysis of monophosphate to nucleoside) ^107^. Both *XDH/XOD* and *5NTD* were downregulated in SHO, suggesting that purine catabolism may be suppressed in SHO, consequently promoting biosynthesis of purines-related molecules, such as energy-yielding metabolites to support growth.

#### 3.5.2. DEUs associated with metabolites and solute transport

The DE analysis also detected four key DEUs involved in cellular solute movement. One of these transporter genes is nose resistant to fluoxetine protein 6-like (*Nrf6*), a transmembrane protein involved in the transport or modification of xenobiotic compounds or particular lipids ^108^. In addition, three members of solute carrier family were identified, namely, sodium-coupled monocarboxylate transporter 1 (*SLC5A8*), solute carrier family 28 member 3 (*SLC28A3*), and solute carrier family 22 member 15 (*SLC22A5*). *SLC5A8* encodes for a Na^+^/glucose co-transporter that facilitates in the transport of monocarboxylates, including short-chain FAs and nicotinate ^109,110^, *SLC28A3* encodes for pyrimidine and purine nucleosides transporter ^111^, and *SLC22A5* encodes for an organic cation transporter and carnitine symporter ^112^ to facilitate carnitine-mediated transport of long-chain FAs from the cytosol to mitochondria for subsequent beta-oxidation and energy production ^113^. We also detected the solute transporter genes *SLC23A1, SLC26A10*, and organic cation transporter-like (*Orct*) in two hatchery datasets. *SLC23A1, SLC26A10*, and *Orct* were upregulated in SHO, suggesting a higher influx of their target molecules (e.g. carnitine, nucleosides, FAs, and cations) to their respective sites of metabolism to induce cellular activities, such as signaling activation, metabolite biosynthesis, and xenobiotic metabolism, consequently influencing the growth of juvenile sea cucumber.

#### 3.5.3. DEUs associated with fatty acid metabolism

The DEU analysis identified three key unigenes that encode for enoyl-CoA delta isomerase 1, cytochrome P450 4V2, and FA binding protein 3. These genes are involved in the mitochondrial fatty acid (FA) beta-oxidation, which plays a pivotal role in energy derivation through degradation of FAs ^114,115^. Upregulation of these unigenes suggests that SHO individuals have higher FA metabolism and mobilization compared to STU, possibly for activating FA-mediated cell signaling pathways or energy production directed for growth.

#### 3.5.4. Death domain-containing protein, branched-chain amino acid aminotransferase-like, and proline-rich transmembrane protein 1

Unigenes identical to death domain-containing protein 1 (*DTHD1*), branched-chain amino acid aminotransferase-like (*BCAT*), and proline-rich transmembrane protein 1 (*PRRT1*) were also identified as key DEUs. Information on *DTHD1* function is lacking ^116^; however, it has been suggested to be involved in activation of apoptosis and inflammatory signaling transduction, which is consistent with the known functions of proteins in the death domain superfamily ^117^. *BCAT* is involved in the catabolism of branched-chain amino acids (e.g. leucine, isoleucine, and valine), generating alpha-ketoacids and glutamate in the process ^118^. Glutamate is a precursor molecule for the biosynthesis of various biomolecules including amino acids (proline and arginine), neurotransmitters (e.g. gamma-aminobutyrate), and glutathione ^119^, while alpha-ketoacids may be further catabolized by other enzymes to final products (e.g. acetyl-CoA) that are consumed in tricarboxylic acid (TCA) cycle to promote fatty acid oxidation and energy production ^120^. Therefore, BCAT may play a role in growth and development through regulation of branched-chain amino acids, glutamate, and alpha-ketoacids-mediated FA metabolism. *PRRT1* has been shown to influence synapse development and function by regulating AMPA receptors in the brain ^121^. It is possible that PRRT1 also participates in the development of nervous system in sandfish.

### 3.6. *In silico* microsatellite and SNP markers discovery

Variant discovery was performed on the *H. scabra de novo* transcriptome to mine potential microsatellite and SNP markers. A total of 47,127 microsatellites, distributed across 35,914 unigenes, were recovered from the final assembly (Table 4 and Additional File 1 Table S10). Of these, 8,422 unigenes contained more than one microsatellite. Mononucleotide motif dominated the microsatellite types accounting for 86.6% of the total repeat motifs, followed by dinucleotide constituting 8.2%.

**Table 4.** Statistics of microsatellite and SNP identified from juvenile *H. scabra* transcriptome.

| Variant type | Classification | Count (%) |
| --- | --- | --- |
| **Microsatellite** | **Total** | **47,127 (100)** |
|  | Mononucleotide | 40,819 (86.6) |
|  | Dinucleotide | 3,875 (8.2) |
|  | Trinucleotide | 1,441 (3.1) |
|  | Tetranucleotide | 845 (1.8) |
|  | Pentanucleotide | 124 (0.3) |
|  | Hexanucleotide | 23 (0.05) |
| **SNP** | **Total** | **373,196 (100)** |
|  | Non-CDS | 310,678 (86.2) |
|  | In CDS | 62,380 (16.7) |
|  | Transitions | 226,265 (60.6) |
|  | A/G | 114,304 (30.6) |
|  | C/T | 111,961 (30) |
|  | Transversions | 146,931 (39.4) |
|  | T/A | 52,981 (14.2) |
|  | A/C | 35,645 (9.5) |
|  | G/T | 33,531 (9) |
|  | C/G | 24,774 (6.6) |

The KisSplice pipeline discovered 373,196 SNPs, which were distributed to 52,729 unigenes. Of these, 86.2% were not in coding sequence (non-CDS), while SNPs detected in coding region (16.7%) comprised 37,191 synonymous and 25,189 non-synonymous types (Table 5). There were more transitions (60.6%) compared to transversions (39.4%) among the final SNP sets. SNP markers developed from the transcriptome have added value because they can be used to study selection and local adaptation to different environmental conditions at spatial and temporal scales ^122^. Therefore, the gene-associated SNPs derived from *H. scabra* transcriptome may be valuable in population genomics studies, especially when loci under selection (non-neutral) that have a direct functional impact are of interest.

KissDE identified 10,959 potentially growth category-associated SNP (p-adjusted cut-off of < 0.01) (Additional File 1 Table S11). Further filtering growth category-associated SNPs with |Deltaf/DeltaPSI| ≥ 0.5 as threshold reduced the number to only 91 SNPs with high potential of being specific to a growth category. The absolute value of Deltaf/DeltaPSI, a KissDE statistic based on allele frequency differences between two conditions, ranges from 0 to 1, in which a SNP with a value of 1 suggests the SNP has a high probability of being condition specific and could present as a fixed allele for a particular condition ^55^. A separate investigation will be necessary to genotype these SNPs and evaluate their utility to differentiate SHO and STU, particularly in view of the pooled sequencing strategy used here which may affect the accuracy of allele frequency estimates used in KissDE. Nonetheless, these putative SNPs represent potential molecular markers to enable marker-assisted selection programs for enhanced growth rates in sandfish.

### 3.7. Comparison with previous studies investigating growth variation in sea cucumbers

Previous transcriptome analysis of growth variation in sea cucumbers have been limited to *A. japonicus*. Downregulation of immune-related genes in slow-growing individuals was associated with global hypometabolism ^123,124^, a physiological state similar to hibernation to cope with stress due to unfavorable conditions. Similarly, a recent transcriptome study on growth of two populations of *A. japonicus* and their hybrid has also highlighted the overexpression of defense- and immune-related genes, such as heat shock protein (HSPs) genes, in slow-growing individuals ^26^. In contrast, our results reveal immune response activation in the fast-growing group based on higher population-wide expression of DEUs possibly encoding for immunity and defense-related genes. Contrasting gene expression patterns between *A. japonicus* and *H. scabra* were also observed for several genes involved in different metabolic processes, including serine protease, *PCSK9*, and IgGFc-binding protein. Further, several key genes reported to be directly associated with growth and development in *A. japonicus* were not detected in *H. scabra*, such as ribosomal proteins (RPLs) and growth factors. While differences in expression patterns of some genes were observed, we also found similar genes with concordant expression in fast-growing individuals for both species, such as fibropellin, *ECI1, SLC28A3, Orct*, and *DHRS4*.

With the work presented here, genes implicated in immune response, solute transport, and energy metabolism are likely involved in growth variation observed in early juvenile *H. scabra* as evidenced by the concordant patterns of expression (key DEUs) observed across three different hatchery populations. Contrasting results for *A. japonicus* and *H. scabra* indicate that genomic mechanisms underlying growth regulation are complex and varies among different sea cucumber species. Consequently, it is imperative to determine the detailed roles of the differentially expressed genes identified in both species to gain further insights on growth variation in sea cucumbers.

## 4. Conclusions

This research presented a *de novo* assembly of the early-stage juvenile *H. scabra* transcriptome and identified genes that are potentially associated with growth variation in juvenile sandfish. DEUs between fast- and slow-growing juvenile sandfish across three hatchery populations were related to potentially key molecular pathways and biological processes controlling growth variation, which include carbohydrate binding, ECM organization, fatty-acid metabolism, and metabolite and solute transport. DEUs related to immunity and defense and energy metabolism were upregulated in fast-growing juvenile sandfish, suggesting that they possess a more robust pathogen-defense response and a higher energy output to sustain increased growth rate. Our results also revealed a large number of potential microsatellites and growth category-associated SNP markers. Functional studies on these genes and SNPs are required to elucidate their roles in growth regulation in sea cucumbers. Overall, our findings improve the current understanding on the genetic basis of growth variation in sea cucumbers and represents an invaluable genomic resource to facilitate future functional genomics-based research and applications in sandfish and other sea cucumbers, including selecting for genes associated with faster-growing phenotypes for marker-assisted selection and broodstock enhancement.

## 5. Availability of data and materials

All raw Illumina data were submitted to NCBI Short Read Archive (SRA) Sequence Database (Bio-Project: PRJNA433757); Accession Numbers: SRR6714451 – SRR6714458 and SRR8713066 – SRR8713073). The final assembly used in all subsequent analyses is available in NCBI’s Transcriptome Shotgun Assembly database under the TSA accession GIRH01000000. Additional File 3 contains the annotation result of Trinotate and diamondblast.

## 6. Acknowledgments

The authors would like to thank the following people and institutions for providing samples and facilitating their collection: D. Ticao of Alson Aquaculture Corp.; M.A. Meñez, J.R. Gorospe, C. Edullantes, B. Rodriguez, A. Rioja, T. Catbagan, and G. Peralta of Bolinao Marine Laboratory, UP-MSI; and E. Tec of Palawan Aquaculture Corp. We also thank K.T. Gulay for providing valuable logistical support for the collection and processing of samples for sequencing.

## 7. Author’s contribution

**JFFO**: Conceptualization, Methodology, Formal analysis, Investigation, Data curation, Visualization, Writing – Original Draft, Writing – Review & Editing, Project administration; **GGG**: Conceptualization, Methodology, Formal analysis, Investigation, Data curation, Writing – Original Draft, Writing – Review & Editing, Project administration; **RRG**: Conceptualization, Methodology, Supervision, Funding acquisition, Writing – Review & Editing, Project administration;

## 8. Conflicts of interest

The authors declare no conflict of interest.

## 9. Funding

This work was supported by the Department of Science and Technology – Philippine Council of Agriculture and Aquaculture Resources Department.

## Notes

### Competing Interest Statement

The authors have declared no competing interest.

