## Additional File 2 for "Transcriptome analysis of growth variation in early juvenile stage sandfish *Holothuria scabra*"

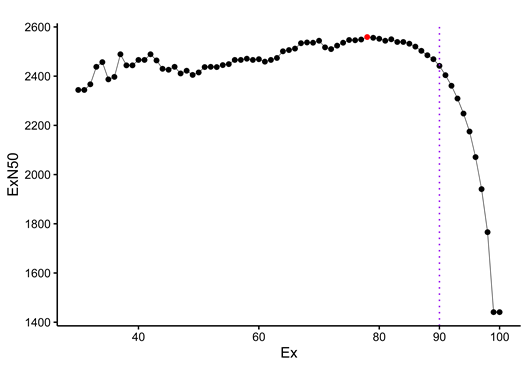

**Figure S1.** ExN50 peak saturation graph of the juvenile stage *H. scabra* final assembly representing the percentage of the total normalized expression data (x-axis) with the relative N50 values (y-axis). Highest point (red dot) is detected at E78.

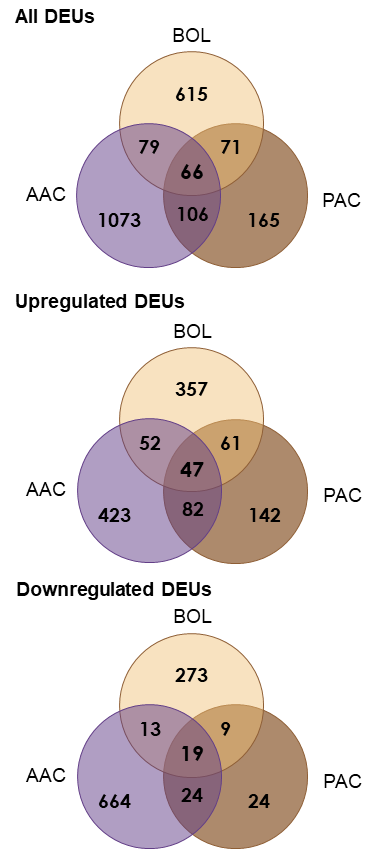

**Figure S2.** Venn diagram representation of significant DEUs (log2FC| > 2, FDR < 0.01) common between and unique in AAC, BOL, and PAC dataset.
